# Regeneration of *Lumbriculus variegatus* requires post-amputation production of reactive oxygen species

**DOI:** 10.1101/2024.06.10.598261

**Authors:** Freya R. Beinart, Kathy Gillen

**Affiliations:** Kenyon College, Molecular Biology, Gambier OH USA; Kenyon College, Biology, Gambier OH USA

**Author notes:** Corresponding Author: Kenyon College 202 N. College Rd. Gambier OH 43022.

**Keywords:** animals, annelids, reactive oxygen species, regeneration, signaling

## Abstract

Animals vary in their ability to replace body parts lost to injury, a phenomenon known as restorative regeneration. Uncovering conserved signaling steps required for regeneration may aid regenerative medicine. Reactive oxygen species (ROS) are necessary for proper regeneration in species across a wide range of taxa, but it is unknown whether ROS are essential for annelid regeneration. Since annelids are a widely used and excellent model for regeneration, we sought to determine whether ROS play a role in the regeneration of the highly regenerative annelid, *Lumbriculus variegatus*. Using a ROS-sensitive fluorescent probe we observed ROS accumulation at the wound site within 15 minutes after amputation; this ROS burst lessened by 6 hours post amputation. Chemical inhibition of this ROS burst delayed regeneration, an impairment that was partially rescued with exogenous ROS. Our results suggest that similar to other animals, annelid regeneration depends upon ROS signaling, implying a phylogenetically ancient requirement for ROS in regeneration.

## Introduction

Regeneration, the remarkable biological phenomenon allowing organisms to replace or repair damaged or lost tissues, occurs in a wide spectrum of capabilities across the animal kingdom. The extent to which animals can regenerate generally decreases in more complex organisms, and may vary through developmental stages or anatomical locations within a species (Zhao et al., 2016). For example, vertebrates, especially mammals, typically exhibit limited regenerative abilities. They can, to some extent, repair and regrow certain tissues and organs, like skin, bone, and the liver, but are generally unable to regenerate complex structures, such as limbs or body parts (Iismaa et al., 2018). Some phylogenetically more basal vertebrates, such as salamanders and zebrafish, exhibit a greater regenerative potential compared to that found in mammals, with the capacity to regenerate amputated limbs and fins, respectively (Akimenko et al., 2003; Joven et al., 2019). In contrast, many invertebrates have remarkable regenerative abilities that far surpass those of vertebrates. This striking capacity for regeneration extends to the regrowth of entire organisms from small fragments with astonishing precision and complete functional restoration. *Hydra* can regenerate missing body parts upon either transverse or longitudinal amputation (Reddy et al., 2019), while the flatworm *Planaria* can regenerate new heads, tails, sides, or entire organisms from small body fragments (Reddien, 2018).

Annelids, a large phylum of segmented worms, vary widely in their regeneration ability (Kostyuchenko and Kozin, 2021), making them an excellent model for comparative studies. Annelids regenerate primarily through epimorphosis, a mechanism involving the formation of a blastema, a mass of undifferentiated cells, which subsequently undergoes extensive proliferation and differentiation to restore lost tissues (Kostyuchenko and Kozin, 2021). This process of restorative regeneration is not solely triggered by injury; many annelids undergo asexual reproduction by self-amputation (autotomy), caused by muscular contraction (Bely, 2014). Following segment amputation via autotomy or injury, the epithelial tissues immediately close around the wound via the rapid constriction of circular muscles at the site of amputation (Kostyuchenko and Kozin, 2021). After the healing of the wound, cells migrate to the wound site and form the blastema (Herlant-Meewis, 1964; Nikanorova et al., 2020; Zattara et al., 2016). Cells in the blastema begin to proliferate and differentiate to form different structures: a prostomium at the anterior end and a pygidium at the posterior end (Martinez Acosta et al., 2021). At the distal end of the pygidium, proliferating cells contribute to the re-establishment of a posterior growth zone, with segmentation occurring via similar mechanisms to those during ordinary posterior segment addition (Martinez Acosta et al., 2021; Özpolat and Bely, 2016). For segmentation of the anterior regenerate, patterning occurs at the blastemal mass, organizing cells into discrete polarized arrangements (Martinez Acosta et al., 2021). After cell differentiation completes, growth allows for the regenerated structure to regain its original proportions and function.

Previous work suggests the most recent common ancestor of annelids was able to regenerate both anterior and posterior ends, but subsequent evolution led to the loss of this ability in various lineages, resulting in a spectrum of regenerative diversity (Zattara and Bely, 2016). Some annelid species in families such as Sabellidae, Chaetopteridae, and Lumbriculidae exhibit extensive regeneration with the ability to regenerate an entire organism from a single mid-body or trunk segment (Bely, 2006). The ability to regenerate fully from trunk segments has been lost in some clades, like the Capitellidae species, *Capitella teleta,* and the Syllidae species, *Amblyosyllis formosa*, where anterior regeneration of the head segments cannot be completed, but extensive segment regeneration is still possible posteriorly (Ribeiro et al., 2018; Seaver and de Jong, 2021). In contrast, the Hirudinea leeches have very limited regenerative capabilities, with the complete loss of axial regeneration capability unlike other closely related annelids (Kuo and Lai, 2019). For organisms like leeches which have lost the capability of regeneration, wound healing is still observed, but they do not form a dedifferentiated blastema that can give rise to many different cell types (Huguet and Molinas, 1994; Odelberg, 2004). This diversity within the Annelida phylum allows for comparative studies between evolutionarily proximal and morphologically similar organisms that can reveal insights into the evolution of regenerative processes.

Understanding the molecular mechanisms that underlie the phylogenetic changes in regenerative ability has emerged as a prominent challenge in the fields of developmental and regenerative biology. There is a heavily skewed focus on a narrow number of regenerative research organisms, limiting what can be learned about regeneration (Sánchez Alvarado, 2018). Introducing new organisms into regenerative research is critical to the field, as a comprehensive examination of signaling across diverse regenerative species can uncover shared mechanisms (Seifert et al., 2023). Studying the mechanisms underlying regeneration in annelids allows a deeper understanding of the processes that result in the variation of regenerative capability.

Although the specific processes may vary among regenerative models, the role of reactive oxygen species (ROS) as a key contributor to the orchestration of regenerative processes is becoming increasingly apparent (Bideau et al., 2021). ROS are highly reactive products of oxygen metabolism. In animals, major sources of cellular ROS are the mitochondrial electron transport chain and membrane NADPH oxidases (NOX), which generate products with relatively higher reactivities than molecular oxygen (O_2_) (Niethammer, 2016; Palma et al., 2024; Thannickal and Fanburg, 2000). These partially reduced metabolites include superoxide (O_2_•–) and hydrogen peroxide (H_2_O_2_), and are detoxified by antioxidant enzymes such as catalase and superoxide dismutase to protect against oxidative damage (Lü et al., 2010; Thannickal and Fanburg, 2000). While often discussed in their role in oxidative stress, it has become increasingly clear that ROS are not purely toxic by-products of cellular metabolism, but instead in regulated concentrations act as key components of cell signaling. Tightly regulated by cellular redox mechanisms, ROS interact with signaling molecules critical for a variety of necessary cellular processes such as proliferation, survival, and apoptosis (Bardaweel et al., 2018; Ray et al., 2012).

The production of ROS in response to injury is highly conserved in animals (Suzuki and Mittler, 2012). By accumulating at wound sites, ROS aid in wound healing by acting as second messengers, or by directly interacting with thiol groups to regulate downstream protein activity (Huizen et al., 2022; Niethammer, 2016). Although regeneration closely follows wound healing, variation in the accumulation of ROS between regenerative and non-regenerative processes indicates that these mechanisms are likely distinct (Owlarn et al., 2017). In regenerative animals, the accumulation of ROS at the wound site is both greater in concentration and sustained for a greater period of time than during non-regenerative wound healing processes (Gauron et al., 2013; Huizen et al., 2022). While in many animal models ROS accumulation is required for successful regeneration (Carbonell M et al., 2022; Gauron et al., 2013; Love et al., 2013; Pirotte et al., 2015), the involvement of ROS in the regeneration of annelids has not yet been explored.

Understanding the potential involvement of ROS in annelid regeneration carries significance for using them as model organisms to uncover regenerative processes applicable to other animals, such as vertebrates. *Lumbriculus variegatus*, our model annelid, is among the most highly regenerative annelids, capable of undergoing complete regeneration of lost body segments (Martinez Acosta et al., 2021). Given the role of ROS in the regeneration of other animals, we hypothesized that regeneration in *Lumbriculus* also depends upon ROS production. To test our hypothesis, we investigated the necessity of ROS accumulation for axial regeneration of *Lumbriculus* using two approaches: first, we characterized the post-amputation spatial and temporal accumulation of ROS in *Lumbriculus*; and second, we investigated the functional consequences of ROS inhibition after the amputation of *Lumbriculus*. We predicted that ROS would accumulate at the wound site of injured *Lumbriculus*, and that inhibiting ROS production would impair regeneration. Our observations provide evidence supporting the role of ROS as a critical signaling mediator in annelid regeneration.

## Materials and Methods

### Animal care and handling

*Lumbriculus variegatus* were purchased from Carolina Biological Supply (Burlington, NC). As there are two clades of *Lumbriculus* which may merit distinct species status (Martinez Acosta et al., 2021), we previously determined that these worms are Clade I (Fischer et al., 2022). Worms were kept in either dechlorinated tap water or artificial freshwater (AFW) containing 0.5g/L Instant Ocean aquarium salts (United Pet Group Inc., Cincinnati, OH, USA) as previously suggested (Crisp et al., 2010) at 16-18°C on a 12 L:D cycle. Worms were fed sinking fish food pellets (Pisces Pros, Ogden, UT, USA) twice each week and water was changed weekly.

Worms were starved at least 48 hours prior to all experiments to allow gut contents to be purged. Only fully grown worms were used for experiments (recently autotomized worms were excluded). Survival of worms during experiments was assessed via brightfield microscopy and gentle probing of the worm segments. For time-course ROS detection experiments, approximately 10 segments in length were amputated from the anterior end and the posterior halves were replaced into DMSO or H_2_DCFDA exposure solutions. For time-course regeneration experiments, worms were briefly placed in 20°C seltzer water for anaesthetization and bisected with microsurgical scissors across the anterior-posterior axis in two locations to produce a trunk fragment; the cuts were made 12 segments and 26 segments from the head. These fragments were photographed at various time points to monitor anterior and posterior regeneration.

### ROS detection assay

General ROS can be visualized *in vivo* using 2′,7′-Dichlorofluorescin diacetate (H_2_DCFDA, Sigma-Aldrich D6883), which is rapidly oxidized to the highly fluorescent 2’,7’-dichlorofluorescein (DCF) in the presence of ROS (Yoon et al., 2018). To assess the validity of using H_2_DCFDA as a fluorescent proxy for *Lumbriculus* ROS levels *in vivo*, endogenous ROS production was induced via heat-stress and blue light irradiation (de Jager et al., 2017; Lockwood et al., 2005). These methods assessed whether the dye could penetrate the cuticle of the worm without amputation-induced cuticle damage. For heat stress experiments, worms were treated in 50μM H_2_DCFDA for 1 hour at 33°C prior to imaging. For blue light irradiation experiments, worms were incubated in 50μM H_2_DCFDA for 1 hour prior to imaging. After incubation, trunks were exposed to 490 nm blue light with an excitation laser for up to 120s and images were taken every 10s to capture DCF fluorescence.

Post-amputation ROS production was visualized *in vivo* using H_2_DCFDA. Stock solutions of 10mM H_2_DCFDA were prepared in DMSO (Sigma-Aldrich D-5879) and kept at −20°C for up to 3 months. Worms were rinsed in AFW and incubated in a solution of 50μM H_2_DCFDA or 0.5% DMSO vehicle control solution for 90 minutes. Worms in exposure solutions remained in the dark prior to imaging. For time-course ROS detection experiments, each worm was imaged a single time at a single time point; worms were not tracked over time. This limits blue light induced ROS production and hence DCF fluorescence.

### Inhibition of ROS production

The general flavoprotein and NOX inhibitor diphenyleneiodonium chloride (DPI, Sigma-Aldrich D2926) was used to inhibit endogenous ROS production in *Lumbriculus.* 3mM DPI stock solutions were prepared in DMSO. Worms were pretreated in solutions of 6μM DPI or 0.2% DMSO vehicle control for 6 hours prior to amputation. After amputation worm fragments were replaced into the exposure solutions (6μM DPI or 0.2% DMSO) for either 30 minutes (Fig. 3) or 6 hours (Fig. 5). Untreated worms in AFW and uncut worms exposed to the same solutions were included as controls for mortality. After the 6 hour post-amputation (hpa) incubation period, worms were removed from the exposure solutions and replaced in AFW for 8 days to track regeneration.

**Figure 1.**
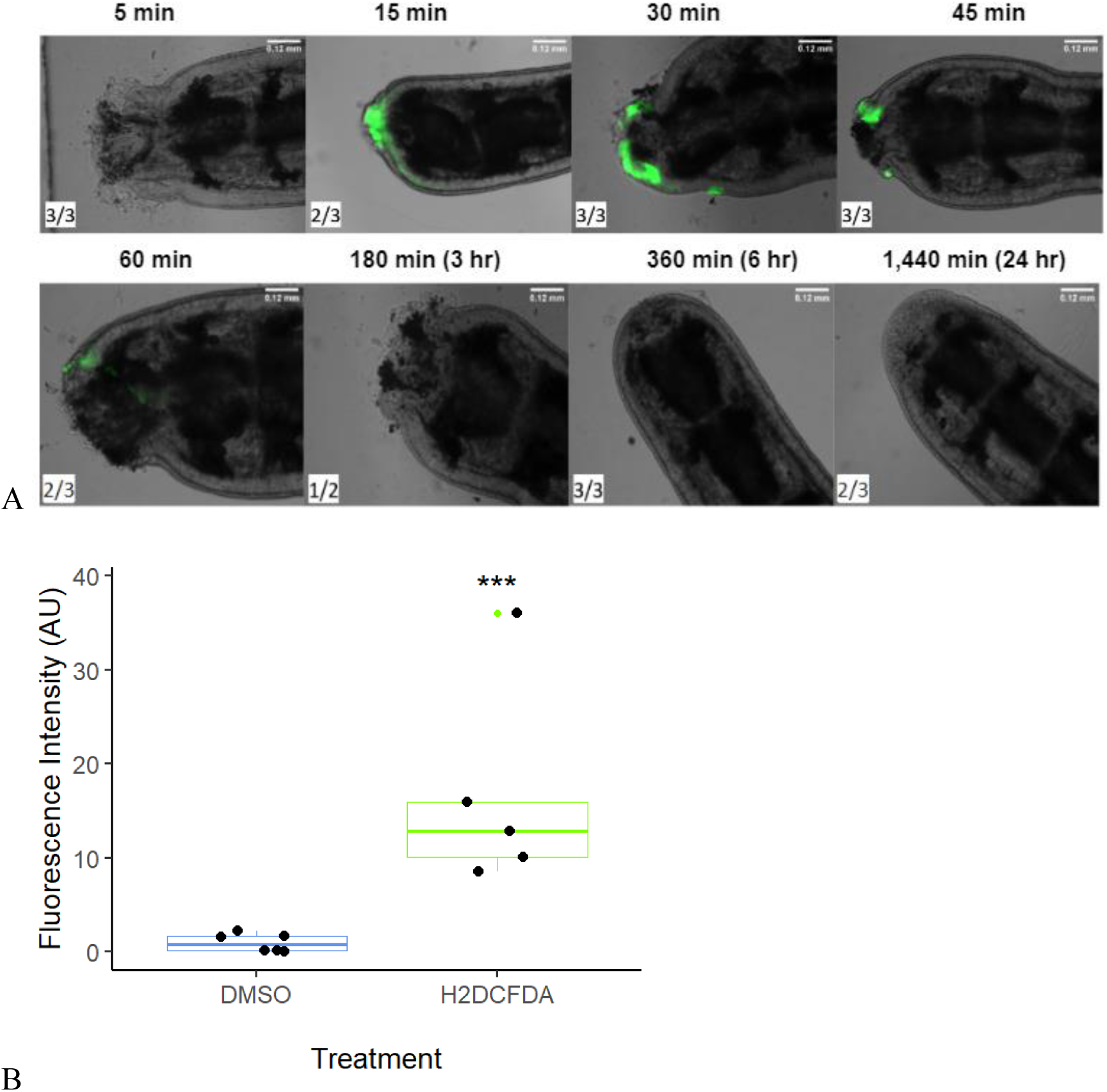
Early increase in ROS production after amputation in *L. variegatus* posterior segments. (A) Worms were incubated in 50uM H_2_DCFDA for 90 minutes prior to imaging at respective intervals of minutes post amputation. For each time point, a representative image is shown. The numerator represents the number of worms displaying a level of fluorescence similar to the representative image and the denominator indicates the sample size of the imaged worms. n= 24 (3 different worms for each time interval). (B) Increased fluorescence intensity of H_2_DCFDA in *Lumbriculus* 30 minutes post-amputation compared to 0.5% DMSO control group. n=6 per treatment group (Welch Two Sample t-test, p = 0.008).

**Figure 2.**
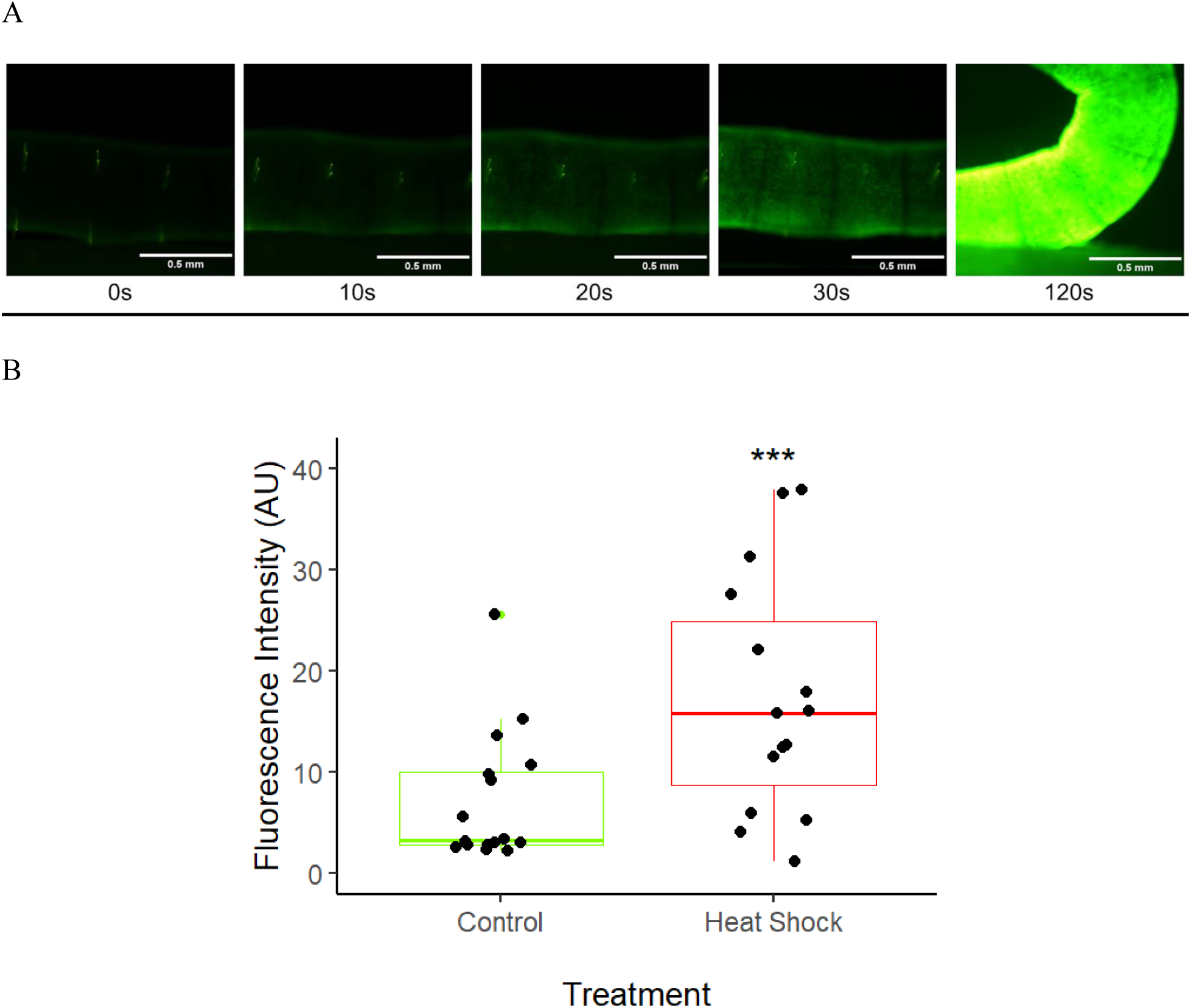
Fluorescence increases during blue light irradiation (A) and heat stress (B) in intact *L. variegatus* treated with H_2_DCFDA. (A) Worms were incubated in 50μM H_2_DCFDA for 60 minutes prior to imaging, then exposed to blue light for a total of 120s. Micrographs show a single worm exposed to blue light for the stated amount of time. n= 5 biological replicates. (B) Worms were incubated in 50μM H_2_DCFDA for 1 hour at 33C or room temperature. n=17 per treatment group (Welch Two Sample t-test, p<0.05).

**Figure 3.**
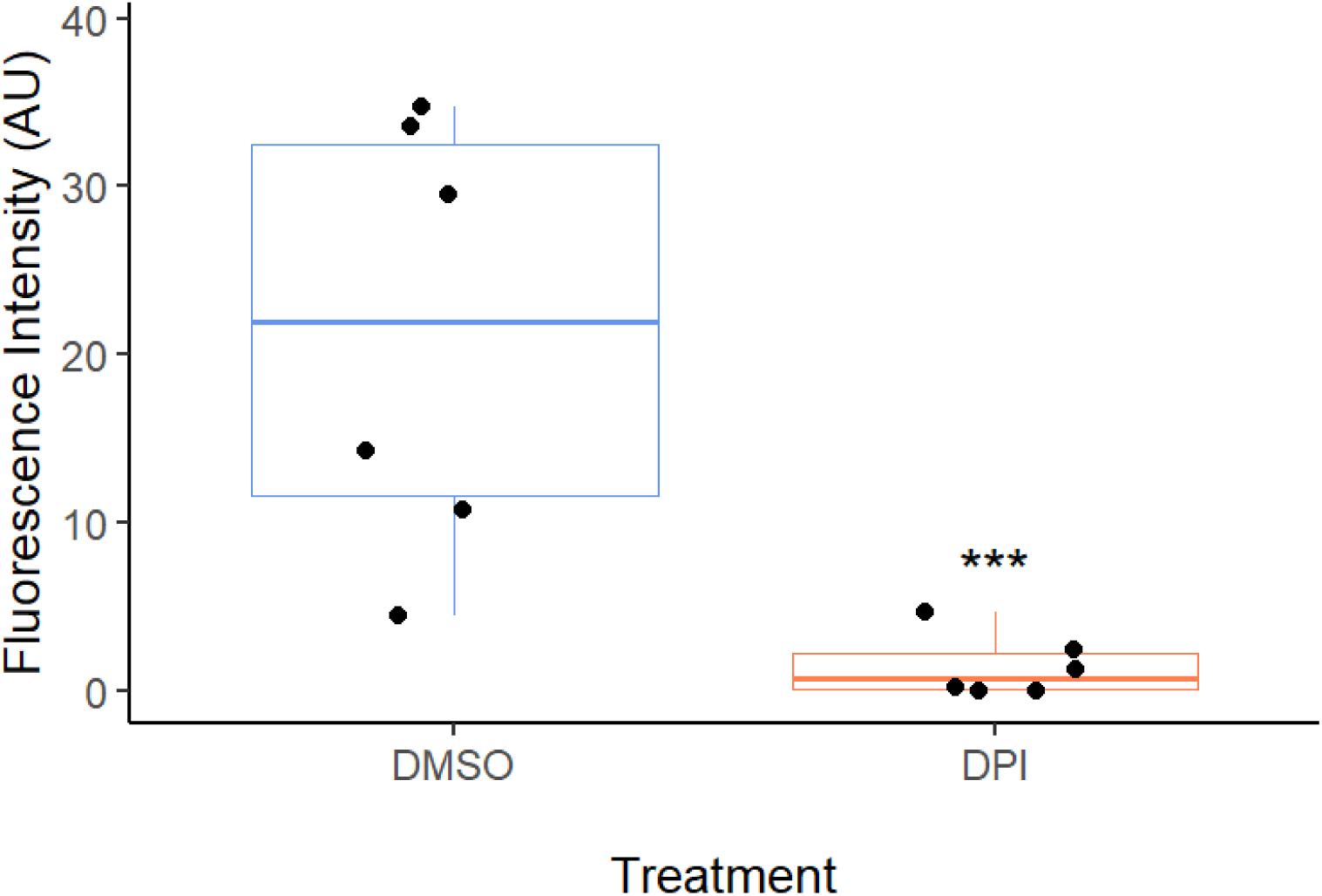
H_2_DCFDA fluorescence intensity decreases in DPI-treated *Lumbriculus* 30 minutes post-amputation. Worms were incubated in a 6μM DPI solution or a 0.2% vehicle control solution for 5 hours prior to addition of H_2_DCFDA. H_2_DCFDA (50μM) was added to the incubation solutions 90 minutes prior to imaging. Worms were amputated ∼10 segments from the head and the posterior segments were imaged 30 minutes post-amputation at the wound site. n=6 per treatment group (Welch Two Sample t-test, p<0.05).

**Figure 4.**
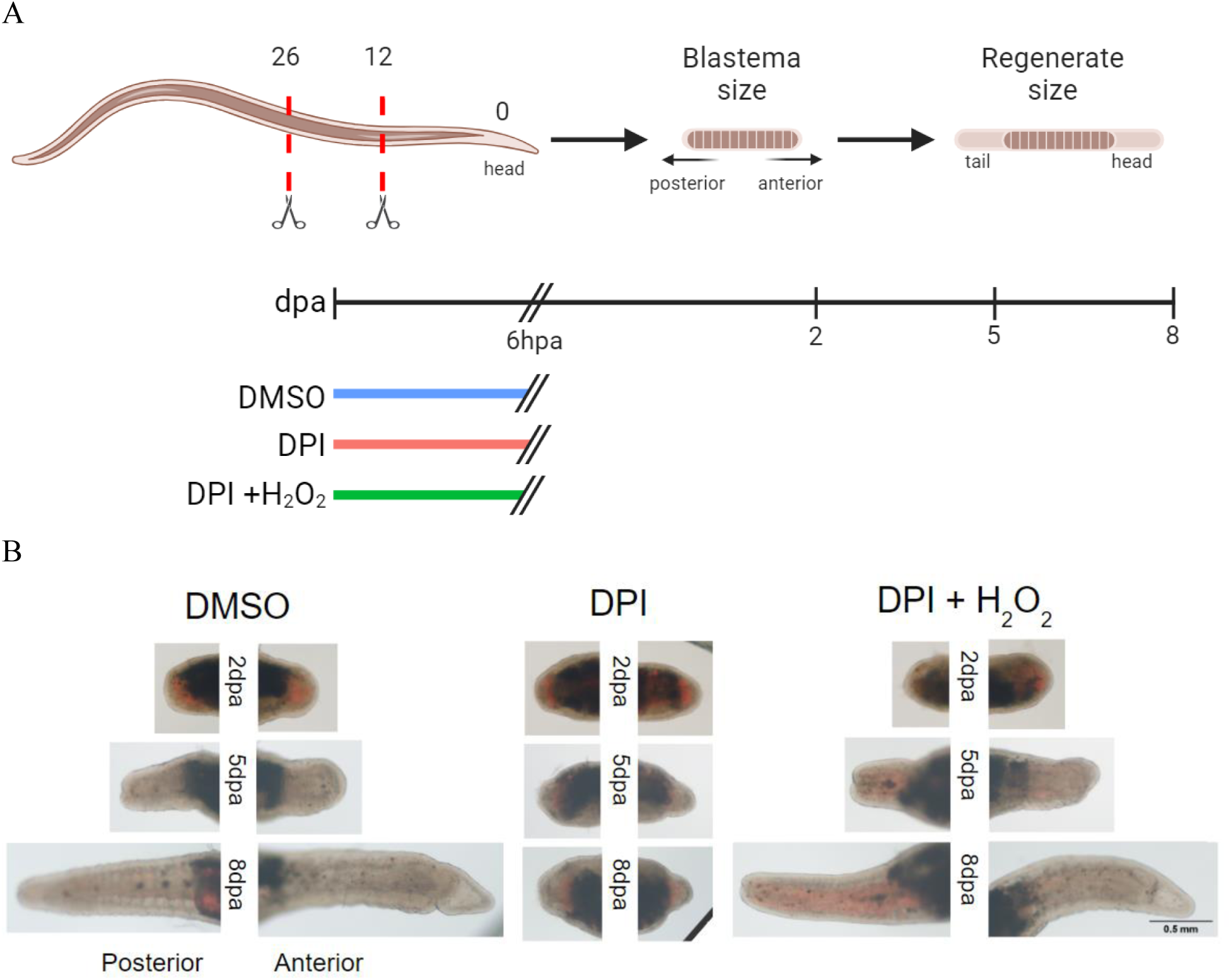
Exogenous ROS rescue regeneration. (A) Schematic of experimental design. Worms were pretreated for 6 h with 6μM DPI or 0.2% DMSO. After amputation, trunk sections were incubated for 6 hours in 0.2% DMSO, 6μM DPI, or 6μM DPI + 50uM H_2_O_2_. Anterior and posterior regeneration was tracked over a period of 8 dpa. (B) Representative images of anterior and posterior regeneration in treated worms. Within each treatment and time point there were no differences in anterior and posterior regenerate area.

**Figure 5.**
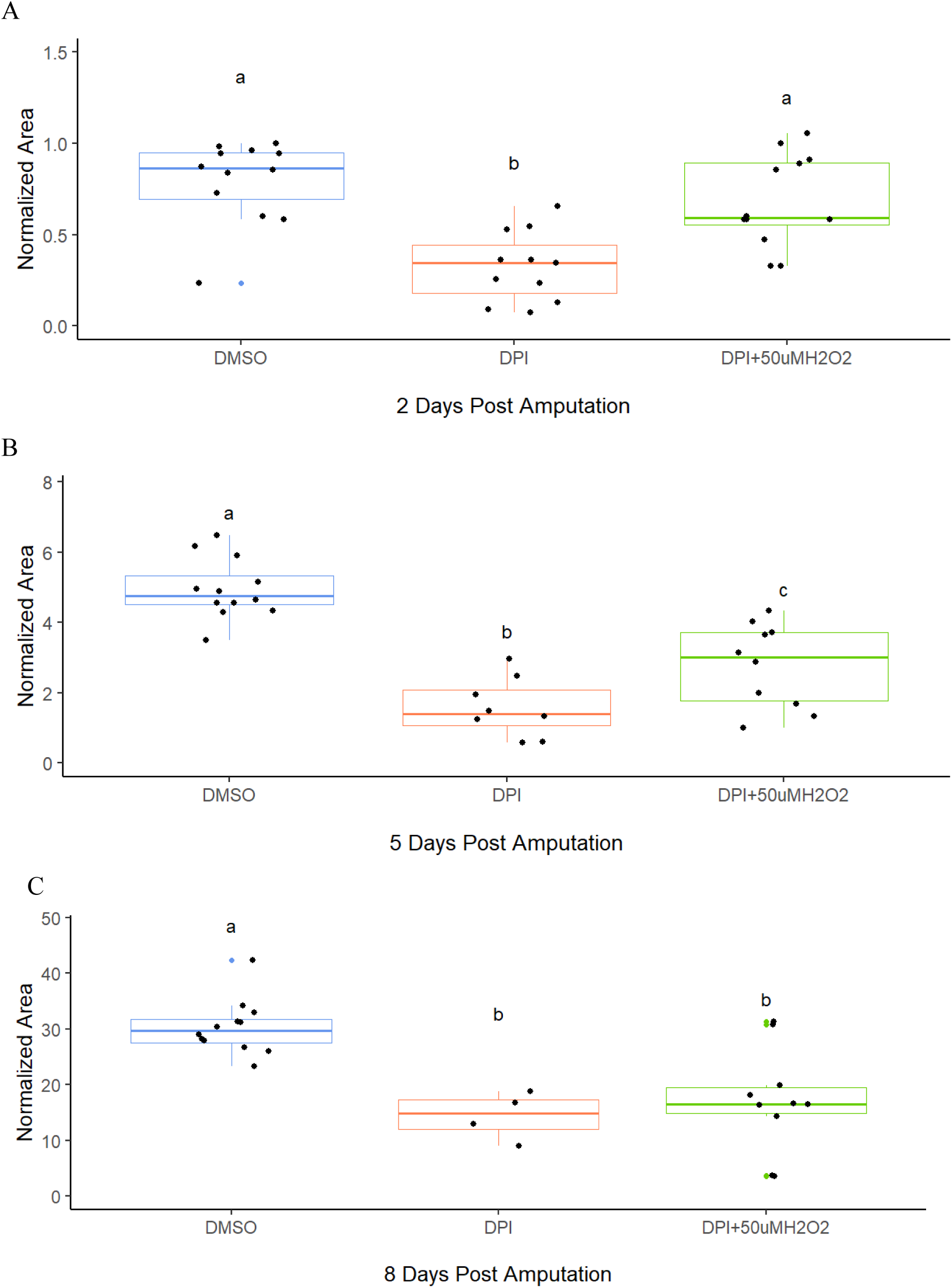
DPI treatment impairs regeneration, and exogenous H_2_O_2_ rescues early regeneration. Amputated worms were treated with 6μM DPI, 6μM DPI supplemented with 50μM H_2_O_2_ or 0.2% DMSO vehicle control for 6 hours post-amputation. Images were captured at **A)** 2 days, **B)** 5 days and **C)** 8 dpa. ImageJ was used to determine areas of new growth based on lack of pigmentation. All areas are normalized to the maximum area of the 2 days post-amputation DMSO worms. Data from both anterior and posterior regeneration is included. Colored dots are outliers, black dots are individual measurements. Sample sizes: (DMSO, DPI, DPI+H_2_O_2_). 2 dpa: (12, 11,12); 5 dpa: (12, 8, 10); 8 dpa: (12, 4, 10). Within each panel, letters represent treatments that were significantly different (one-way ANOVA followed by Tukey HSD post-hoc test, p <0.05).

### Rescuing regeneration using exogenous ROS

All worms were submerged in 6μM DPI or a 0.2% DMSO vehicle control for 6h prior to amputation and trunk sections were replaced in exposure solutions for 6 hpa. For the rescue assay, worms were incubated from 0 hpa to 6 hpa in a 6μM DPI solution containing 50uM H_2_O_2_. Vehicle control worms were incubated in 0.2% DMSO for the same time interval, and a group exposed to the same concentration of H_2_O_2_ in the absence of DPI was included as a control for any H_2_O_2_-related mortality. Posterior and anterior regeneration was tracked using morphological measurements for regenerate structure area over a period of 8 days post-amputation (dpa). If the axial orientation could not be differentiated by morphology, the direction of blood flow in the dorsal vessel was examined, as blood flows from the posterior to the anterior end (Lesiuk and Drewes, 1999).

### Brightfield and fluorescence microscopy

Brightfield and fluorescence images were captured using a Nikon Optiphot-2 microscope equipped with an episcopic fluorescence attachment. For ROS detection experiments, worms were rinsed with dechlorinated water and placed on a glass slide or a 1% agarose pad with a coverslip to minimize movement during live imaging. DCF fluorescence was detected with an excitation wavelength of 490 nm and an emission wavelength of 520 nm. Fluorescent images were taken immediately after light exposure with acquisition settings of 100 ms exposure and 40x or 100x magnification. For regeneration-tracking experiments, worms were rinsed in AFW and placed on a channeled glass slide with a coverslip. Brightfield images were captured at 40x magnification with 1/1000s exposure.

### Data analysis and statistics

Quantification of DCF fluorescence was performed using FIJI ImageJ software (version 2.1.0). The brightness and contrast values of the images were set equally among all data within each experiment. Rectangular regions of interest were selected at the cut site of the worm, with a constant height and a width consistent with the width of each worm. The mean pixel intensities of the green color channel were measured. To assess regeneration, regenerate structure area was measured with thresholding in FIJI ImageJ software (version 2.1.0).

All statistical tests were conducted in R (version 4.2.3). Data were analyzed with either a Welch Two Sample t-test or a one-way ANOVA and Tukey’s HSD post-hoc test, with a significance threshold of p< 0.05. The survival probability of worms was estimated through the non-parametric Kaplan–Meier (log rank) test.

## Results

### Amputation induces ROS production in Lumbriculus

Since ROS are necessary for regeneration in other animals, we hypothesized that ROS are also essential for annelid regeneration. Thus, we expected to observe the production of ROS post-amputation. To assess ROS production, we used the fluorescent probe H_2_DCFDA to measure ROS levels in live worms at several time points post-amputation (Figure 1a). Reactive oxygen species accumulate as soon as 15 minutes after amputation and remain at the wound site until at least 1 hour after amputation. From 6 to 24 hours after amputation, ROS were no longer detected at the amputation site in most worms. One of the worms showed a fluorescence signal localized to the blastema 24 hours after amputation, indicating ROS may be produced past the point of the wound-healing phase of regeneration (Supplementary Figure 1). At 30 minutes post-amputation, worms exposed to H_2_DCFDA exhibited a 15-fold increase in fluorescence compared to vehicle control worms (Figure 1b, t-test, p=0.008), indicating ROS accumulation.

To ensure that the spatial distribution of H_2_DCFDA fluorescence was not due to enhanced penetration at the open wound site, we evaluated its ability to permeate the cuticle by inducing ROS through the punctureless methods of heat and blue light exposure (Figure 2). In both experiments, we found an increase in the H_2_DCFDA signal, indicating the ability of H_2_DCFDA to permeate the worms’ cuticle. The captured fluorescence intensity increased during continuous exposure to blue light (Figure 2a). Blue light itself can convert H_2_DCF into the fluorescent DCF while simultaneously inducing ROS within the organism (Lockwood et al., 2005); however H_2_DCFDA must first be cleaved by cellular esterases. The detected signal was not solely attributed to the blue light cleavage of H_2_DCFDA, as worms treated with H_2_DCFDA exhibited heightened fluorescence intensity in response to heat shock-induced stress even without blue light irradiation (Figure 2b). To eliminate the possibility of the blue light increasing the fluorescence intensity during imaging, all fluorescence microscopy images were taken immediately after exposure to the excitation wavelength.

### Amputation-induced ROS are necessary for regeneration

To explore the role of ROS in regeneration, we sought to decrease ROS levels by inhibiting NOXs, a major source of cellular ROS (Augsburger et al., 2019). We used the flavoprotein inhibitor DPI, a widely used chemical in experiments that aim to decrease endogenous ROS production (Bedard and Krause, 2007; Carbonell M et al., 2022; Gauron et al., 2013; Pirotte et al., 2015). Worms exposed to DPI experienced a 15-fold decrease in H_2_DCFDA fluorescence 30 minutes post-amputation compared to vehicle control worms (Figure 3, t-test, p = 0.01), indicating diminished ROS production. The vehicle control group had a wider range of fluorescence values among individual worms than the DPI treated group. This trend is attributed to the biological variation of endogenous ROS production in individual worms; however, ROS production was decreased to similar levels among all DPI-treated worms.

Having ascertained that DPI could inhibit ROS production, we tested the effects of DPI on axial regeneration. Given that ROS accumulation in most worms decreased by 6 hpa, we exposed worms to DPI until 6 hpa to inhibit the early burst in ROS generation (Figure 4a). The size of both the posterior and anterior regenerate structures at 5 days post-amputation was reduced in DPI-treated worms to approximately a third the size of the vehicle control worms (Figures 4b and 5b). At 8 days post-amputation, the size of new growth in DPI-treated worms was reduced to approximately half the size of the vehicle control worms (Figure 5c). Notably, by day 2 the DPI-treated worms formed unpigmented masses resembling blastemas (Figure 4b), but the growth of these blastemas was stunted, and the regenerate growth never reached the size of the vehicle control worms after this stage of regeneration. Additionally, the survival rate of DPI-treated worms was heavily reduced (log-rank test, p=0.033), with a survival rate of 67% at 5 days post-amputation, and 33% after 8 days (Figure 6).

**Figure 6.**
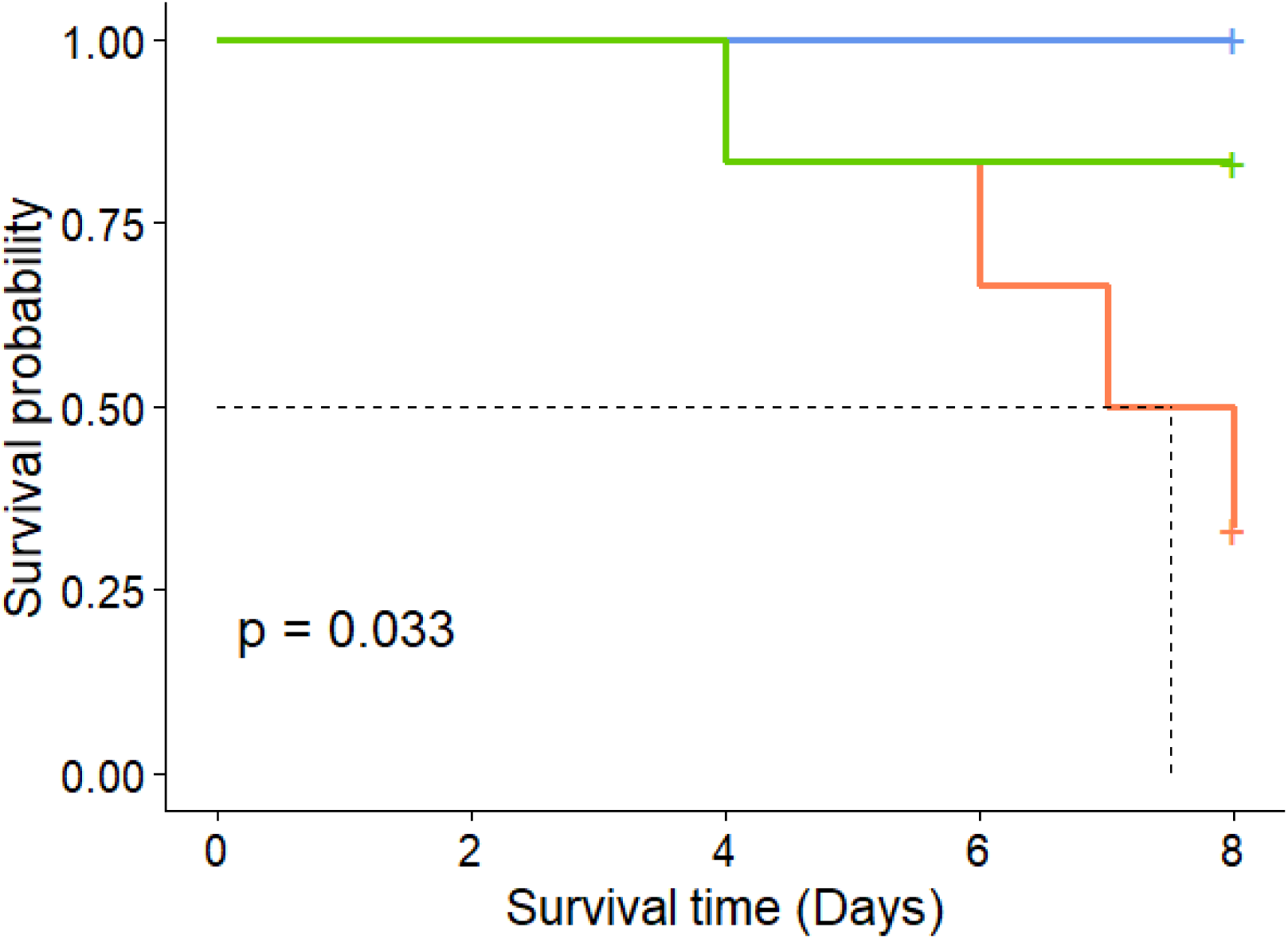
DPI treatment impairs survival, and exogenous H2O2 rescues survival. Amputated worms were treated with 0.2% DMSO (blue), 6μM DPI, 6μM DPI (red), or 6μM DPI supplemented with 50μM H_2_O_2_ (green) for 6 hours post-amputation. The Kaplan–Meier estimates of survival for treatment groups are shown. Significance was determined by the log-rank test, p=0.033, n=6.

### Exogenous ROS rescue survival and regeneration impairments caused by NOX inhibition

We sought to rescue the defects of DPI-treated worms by the addition of exogenous ROS to the media, in the form of H_2_O_2_ (Figure 4a). H_2_O_2_ is a main product of NOX activity, one of the flavoproteins that is inhibited by DPI. The addition of H_2_O_2_ to the DPI media restored the survival rate of the worms to 83% (Figure 6), and trended towards restored regeneration, especially at early time points (Figure 5). At 2 days post-amputation, H_2_O_2_ restored the regeneration of worms (Figure 5a). These restorative effects decreased to a partial rescue of regeneration at 5 days post-amputation (Figure 5b), and did not seem to rescue regeneration at 8 days post-amputation (Figure 5c). Taken together, these results show that post-amputation ROS production contributes to regeneration success and survival of the organism.

## Discussion

We examined the role of ROS in *Lumbriculus* regeneration. ROS accumulated at the wound site within 15 minutes after amputation and diminished in most worms by 6 hours post-amputation. Inhibition of this ROS burst impaired regeneration and decreased survival, the effects of which could be partially rescued with the addition of exogenous ROS. These findings indicate that ROS are crucial for successful annelid regeneration and are consistent with the requirement of ROS for proper regeneration in other animals, including axolotl (Al Haj Baddar et al., 2019; Carbonell M et al., 2022), zebrafish (Gauron et al., 2013), planaria (Huizen et al., 2022), and frog (Love et al., 2013), pointing to a phylogenetically ancient requirement for ROS in regeneration.

While exposing *Lumbriculus* to H_2_O_2_ during the 6 hpa window when DPI was present rescued regeneration at 2 days post-amputation, the H_2_O_2_ rescue effect diminished over time. The inability of our 6 hpa rescue treatment to maintain the regeneration trajectory may point to the necessity of other early produced ROS species for allowing continued growth. Of note, DPI is a general inhibitor of flavoproteins (including nitric oxide synthase) and is not specific to any particular ROS-producing enzyme (Bijnens et al., 2021; Reis et al., 2020; Wind et al., 2010). It is likely that molecules other than H_2_O_2_ are produced post-amputation and contribute to regeneration. Supporting this claim, in planaria DPI caused greater regeneration defects than apocynin (Pirotte et al., 2015), a result the authors attribute to the more specific action of apocynin on NOX enzymes. If reactive species other than H_2_O_2_ are involved in regeneration, this could explain why the DPI-impaired regeneration was not fully restored during H_2_O_2_ rescue. The fluorescent probe H_2_DCFDA lacks specificity for ROS types; using specific probes for particular reactive oxygen and nitrogen species (Kalyanaraman et al., 2012) may pinpoint which are accumulating during *Lumbriculus* wound response and regeneration. It will also be of interest to monitor ROS accumulation for a period of time longer than 24 hours post-amputation, as recent study on axolotls revealed a second ROS burst during blastema formation; the inhibition of ROS during this extended time period resulted in regenerated limbs of reduced size (Carbonell M et al., 2022). In addition, a study in planaria suggests that ROS are needed later than 24 hpa for successful regeneration (Pirotte et al., 2015).

The effects of H_2_O_2_ rescue varied among individual worms. Variability in H_2_O_2_ rescue was also seen in planaria (Jaenen et al., 2021). Planaria fragments were treated with an MEK inhibitor, which prevented regeneration. After washing out the inhibitor, planaria were exposed to H_2_O_2_. In trunk fragments ∼27% of worms underwent normal regeneration via H_2_O_2_ rescue, ∼21% experienced partial regeneration, and ∼45% did not regenerate. Similarly, in tail fragments, ∼34% of the worms underwent full or partial regeneration and ∼52% did not regenerate (14% died). Though the worms used in our experiment were likely genetically clonal and in very similar environments, stochastic gene expression (McAdams and Arkin, 1997) may explain the variability in ROS-related phenotypes. Noisy gene expression of catalases and peroxidases could lead to differences in H_2_O_2_ concentrations in individual cells. Stochastic gene expression within individual cells can lead to differences at the organism level (Raser and O’Shea, 2005), perhaps especially so for a cell permeable molecule such as H_2_O_2_. There is evidence that ROS may act as both upstream activators and downstream effectors of the MAPK pathway (Jaenen et al., 2021); if there is a positive feedback loop, initial random differences can be magnified.

In addition to stochastic gene expression in ROS producing and degrading enzymes, noisy expression of other genes that modulate regeneration may explain biological variation of new growth within each treatment group. Additional variation may arise from our method of measuring new growth. *Lumbriculus* undergo peristaltic movements, where their body stretches and shortens along the posterior-anterior axis. This movement can result in differences in length measurements in a single worm, which is why we chose to measure regenerate area. While area is a more accurate metric of new tissue production than length, volume is even more accurate. A recent study in the annelid *P. leidyi* used a cylindrical formula to approximate volume (Rennolds, 2024). A cylindrical formula would not be appropriate for *Lumbriculus* regeneration as the new growth is conical.

Mechanisms for how ROS promote regeneration are being uncovered. In *Hydra*, the suggested mechanism of ROS signaling in regeneration is via the activation of MAPKs, which, in turn, activate Wnts (Tursch et al., 2022). Likewise, in *Xenopus*, the sustained production of ROS is required for the activation of the Wnt signaling pathway, which promotes cell proliferation and axial growth (Love et al., 2013). Studies in planaria indicate that ROS activate the MAPK/ERK pathway which is required for regeneration (Jaenen et al., 2021). In zebrafish, post-amputation ROS production activates the apoptosis and JNK pathways, both of which aid epidermal cell proliferation (Gauron et al., 2013). The shared element among these pathways is the downstream accumulation of proliferative cells, supporting the idea that cell proliferation is necessary for regeneration. Regeneration also requires the formation of a blastema, which is characterized by proliferative cells acquiring the ability to undergo patterning, differentiation, and growth (Seifert and Muneoka, 2018). It is still unclear if the wound healing process differs among regenerating and non-regenerating animals, as it is unknown whether the specific triggers for blastema formation occur during wound healing or afterwards (Seifert et al., 2023).

In addition to activating signaling pathways necessary for cell proliferation and differentiation (Gauron et al., 2013; Jaenen et al., 2021; Pirotte et al., 2015), ROS directly modulate enzymes that control the cell cycle and the anabolic processes needed to sustain proliferation. For example, ROS oxidation of a cysteine in the cell division cycle 25 protein (Cdc25) modulates its activity and hence the cell cycle; disrupting ROS production caused mitotic arrest via misregulation of Cdc25 (Han et al., 2018). It is becoming clear that regeneration may require elevation of the pentose phosphate pathway (PPP) to provide the metabolites that allow cell division (Patel et al., 2022). ROS-mediated oxidation of a cysteine essential for the catalytic activity of pyruvate kinase M inhibits the enzyme, blocking the last step of glycolysis, and diverting glycolytic intermediates into anabolic pathways such as the PPP (Anastasiou et al., 2011). It remains to be determined if these same pathways and proteins underlie *Lumbriculus* regeneration. If so, it points towards core pathways that may need to be activated to allow tissue regrowth, and suggests clinical targets for promoting therapeutic tissue repair.

While ROS-mediated signaling has been implicated in regeneration of planaria, *Hydra*, *Xenopus*, salamander, and zebrafish models, it has not previously been studied in annelids. Annelids are a phylum with diverse regenerative capabilities; understanding the molecular mechanisms that underlie the differences in the extent to which these organisms can regenerate may aid in unraveling the complexities of tissue repair and regrowth in vertebrates. This study provides the first evidence of ROS involvement in the regeneration of the annelid *Lumbriculus*, opening avenues for further research into ROS signaling and pathways involved in this process. Annelids emerge as a robust model for comparative regenerative studies, suggesting conserved pathways that require ROS for regeneration across animal taxa.

## CREDIT STATEMENT

Freya Beinart: Conceptualization, Methodology, Formal Analysis, Investigation, Writing-Original Draft, Writing-Review & Editing, Visualization Kathy Gillen: Conceptualization, Methodology, Formal Analysis, Resources,Writing-Review & Editing, Supervision

## Acknowledgements

We thank all members of the Gillen lab for their contributions to this project, especially Amanda Caroll, Fielding Fischer, Sydney Buchman, Seryne Rafique, and Emily Banthin. Dr. Kay Tweeten provided method advice using H_2_DCFDA. Dr. Sarah Petersen provided help with fluorescence analysis using ImageJ. Drs. Michael Romero, Iris Levin, Natalie Wright, Joan Slonczewski, Jennifer McMahon and Chris Gillen provided comments on manuscript drafts. Kenyon College provided research funding. We also appreciate the support of the Kenyon College Cascade Program for supporting a summer of research. We used BioRender to create schematic diagrams.

