## Supplementary Figure 1 for "Regeneration of *Lumbriculus variegatus* requires post-amputation production of reactive oxygen species"

**Beinart and Gillen, 2024 Supplementary Figures**

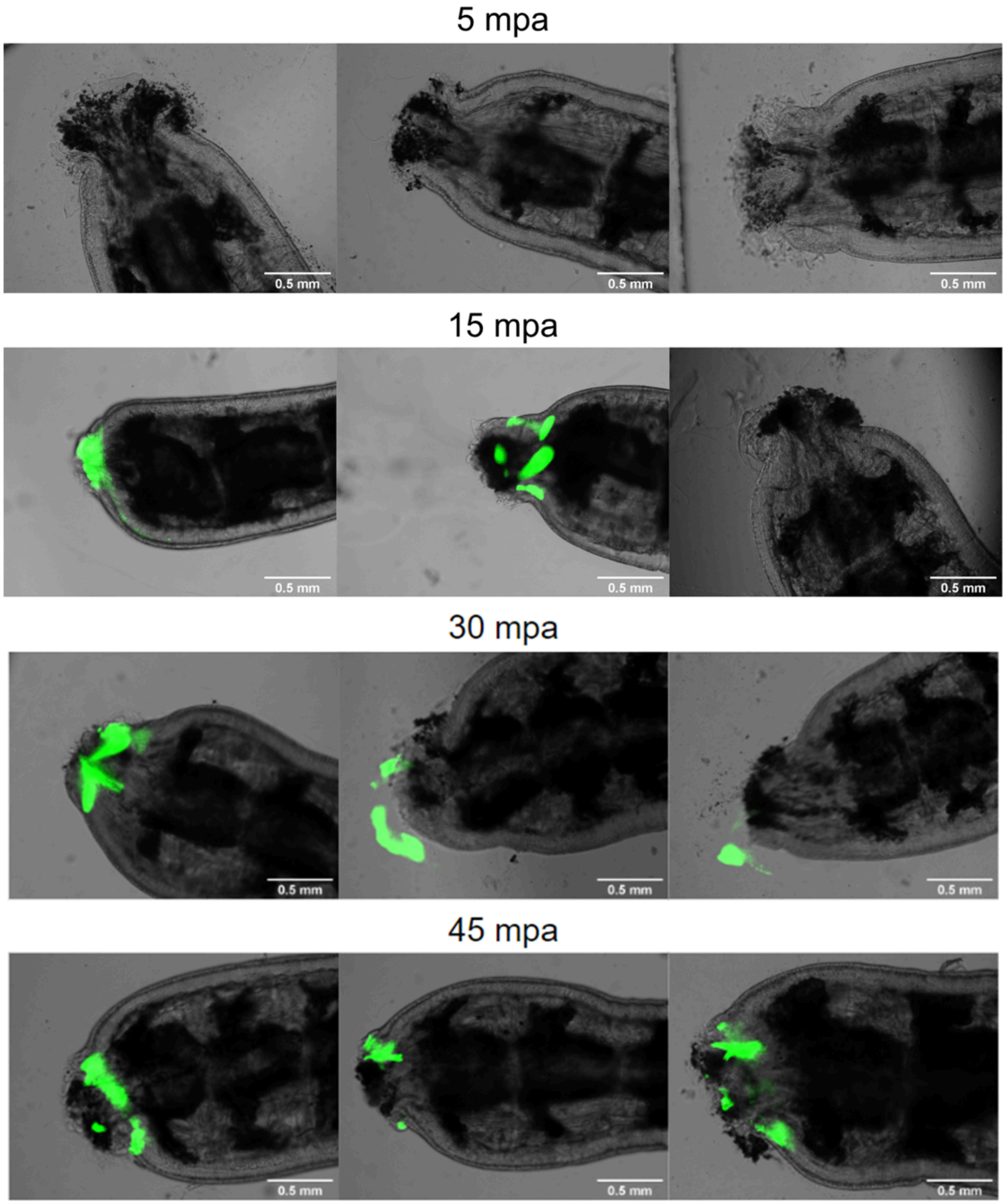

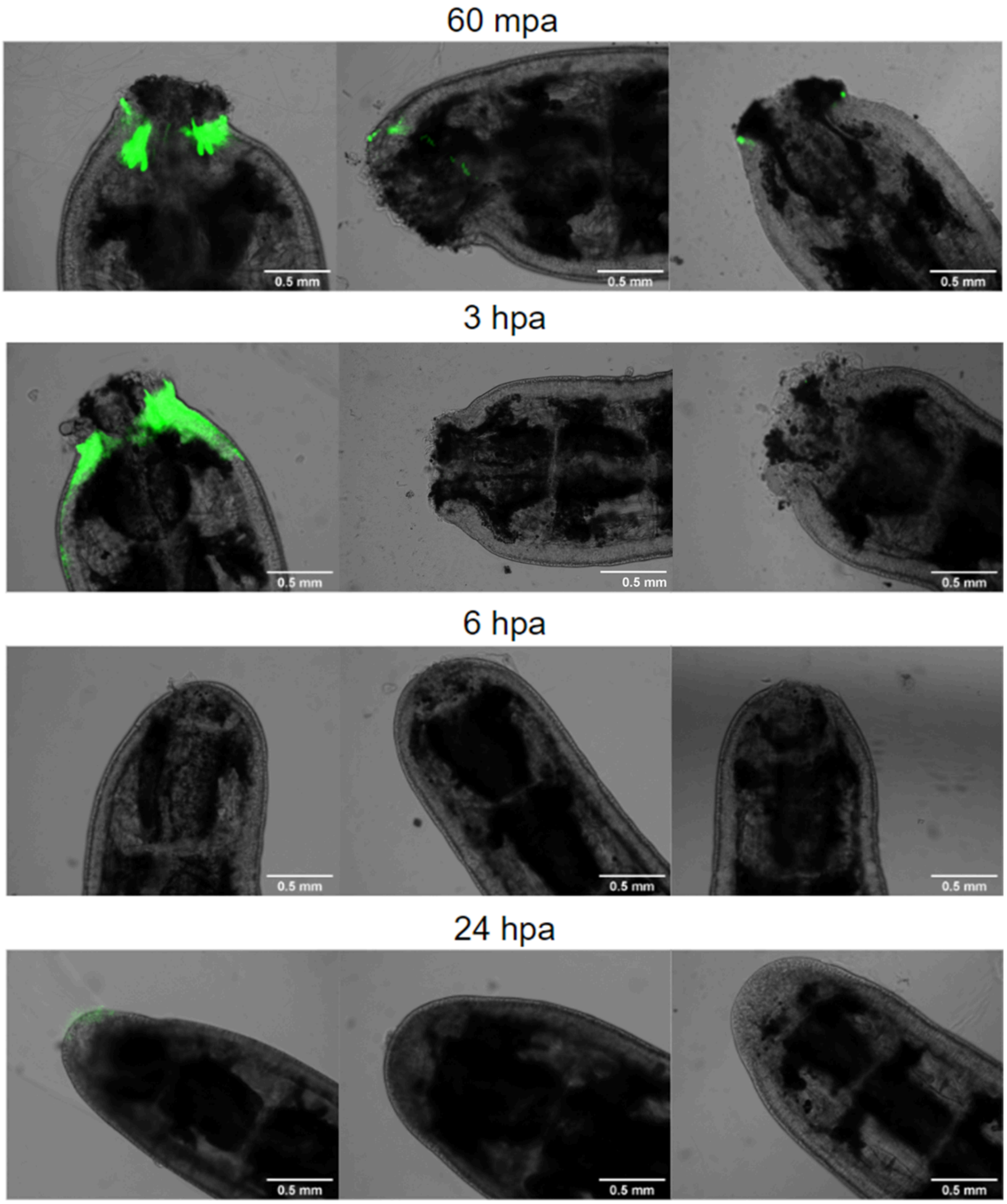


**Figure S1. ROS production after amputation in *L. variegatus* posterior segments.** Worms were incubated in 50uM $H_2$DCFDA for 90 minutes prior to imaging at respective intervals of minutes or hours post amputation. Autofluorescence from chaetae below the wound site was removed in ImageJ. n=3 worms per treatment group.
